# Phenotyping single-cell motility in microfluidic confinement

**DOI:** 10.1101/2021.12.24.474109

**Authors:** Samuel A. Bentley, Vasileios Anagnostidis, Hannah Laeverenz Schlogelhofer, Fabrice Gielen, Kirsty Y. Wan

## Abstract

At all scales, the movement patterns of organisms serve as dynamic read-outs of their behaviour and physiology. We devised a novel droplet microfluidics assay to encapsulate single algal microswimmers inside closed arenas, and comprehensively studied their roaming behaviour subject to a large number of environmental stimuli. We compared two model species, *Chlamydomonas reinhardtii* (freshwater alga, 2 cilia), and *Pyramimonas octopus* (marine alga, 8 cilia), and detailed their highly-stereotyped behaviours and the emergence of a trio of macroscopic swimming states (smooth-forward, quiescent, tumbling or excitable backward). Harnessing ultralong timeseries statistics, we reconstructed the species-dependent reaction network that underlies the choice of locomotor behaviour in these aneural organisms, and discovered the presence of macroscopic non-equilibrium probability fluxes in these active systems. We also revealed for the first time how microswimmer motility changes instantaneously when a chemical is added to their microhabitat, by inducing deterministic fusion between paired droplets - one containing a trapped cell, and the other, a pharmacological agent that perturbs cellular excitability. By coupling single-cell entrapment with unprecedented tracking resolution, speed and duration, our approach offers unique and potent opportunities for diagnostics, drug-screening, and for querying the genetic basis of micro-organismal behaviour.

## 1 Introduction

All lifeforms are environmentally intelligent; even single cells sense and respond to their environment [1]. How living systems interact with the external world is a fundamental question that has fascinated researchers for centuries. The exploratory behaviours of living beings can be extraordinarily complex and context-dependent, and yet also stereotyped. Digitally-assisted and non-invasive tracking methods have revolutionised the study of behaving animals [2, 3]. Algorithms for pose and gait-estimation based on computer vision and machine learning have bolstered the utility of animal models for neuroscience, providing a highly quantitative and objective framework for measuring behaviour. These nascent approaches are now being used in combination with genetic manipulation, to resolve how an organism’s sensory architecture (neural pathways, nervous systems) may be coupled to motor actuators (e.g. muscles) to drive motor actions [4].

Surprisingly, nervous systems are not necessary for the emergence of complex search patterns or behavioural motifs [5, 6]. Micoorganisms also adopt highly stereotyped locomotion strategies, often employing motile appendages: bacterial flagella for swimming, pili for twitching and swarming, and in many eukaryotes, cilia for self-propulsion. Microbial motility is strongly influenced by boundaries, interfaces, and other obstacles [7, 8]. Microorganisms naturally encounter heterogeneous environments, such as soil, sea ice, or porous materials that constitute complex networks of confining tubes, spaces, or interstices. By replicating such microenvironments in the laboratory, realistic situations can be created for tracking and understanding the mechanisms of movement and behavioural adaptation [9, 10].

Despite their small size, microswimmers self-propel at speeds of several tens (even hundreds) of body lengths per second, posing a formidable challenge for live-cell microscopy and tracking. Typically, the same individual can only be observed for relatively short periods, and motility statistics are then coarse-grained across entire populations. Across different species, even across different individuals of the same clonal population, microbes exhibit significantly divergent behaviours. This single-cell heterogeneity may be suppressed if only population averages are considered. However, long-term tracking of the same individual was not possible until very recently [11, 7, 12].

What are the phenotypic or behavioural signatures that distinguish one microswimmer from another? To address this, we studied two species of motile algae, and decoded their distinct responses to environmental cues. The first, *Chlamydomonas reinhardtii* (hereafter CR) is a freshwater biflagellate and a model organism for motility studies and cilia biology [13]. The second, *Pyramimonas octopus* (hereafter PO), is a marine octoflagellate, which has a unique tripartite run-stop-shock motility pattern [14]. Harnessing microfluidics technology to assess long-term cell motility [15, 16], we developed a platform to stably encapsulate algae inside droplets, which are subsequently trapped in quasi-2D microwells. Inside these wells, cells could explore their local environment over much longer timescales than previously possible. We then developed a second apparatus to dynamically perturb the environment of a trapped cell, by droplet pairing and fusion caused by surfactant-induced interface destabilization [17, 18, 19].

We used this approach to phenotype the short and long-term swimming behaviours of the two microalgal species. Long-time tracks allowed us to extract universal metrics for measuring microbial motility, and also led us to the unexpected discovery of non-equilibrium flux loops in the time-averaged motion of single cells in confinement. The effective integration of these complementary experimental and computational techniques (microfluidic trapping, high-speed imaging, single cell tracking, and trajectory analysis) offers an exciting opportunity to understand how living cells interact and respond dynamically to environmental cues (mechanical, light, chemical).

## 2 Results

### 2.1 Microfluidic trapping of single motile algae

We encapsulated CR or PO cells in droplet microwells of four different diameters, spanning a large dynamic range (∅ = 40, 60, 120, 200 *μm*). In order to study the cells’ baseline motility (in the absence of other factors), we used red light illumination to prevent phototactic effects (long-pass IR filter). According to the absorption spectra of CR photoreceptors (ChR1 and ChR2), cells are insensitive to wavelengths of > 600 nm [20]. While the identity of the photoreceptor in PO has not yet been confirmed, it is likely to be similar to CR, since the algal eyespot is highly conserved [21].

Only wells containing precisely one cell were selected for imaging (see Methods). These water-in-oil droplets were unimpeded and retained their shape and volume for more than 1 hour. The trapping procedure was sufficiently gentle and did not damage the cilia or cause spontaneous deciliation. Compared to the cell-scale, the arenas are shallow (~ 30 *μm* deep in total) and quasi-2D, so that swimming motion is largely restricted to the plane. Typical trap dimensions are shown in Figure 1B. We trapped each cell for a total of one hour, splitting the recordings into 5 minute intervals (*N* = 5 cells per trap size, 6 time points each). High-speed imaging was used to resolve rapid motility changes, and cell-centroid trajectories were obtained by particle tracking (see Methods),

**Figure 1:**
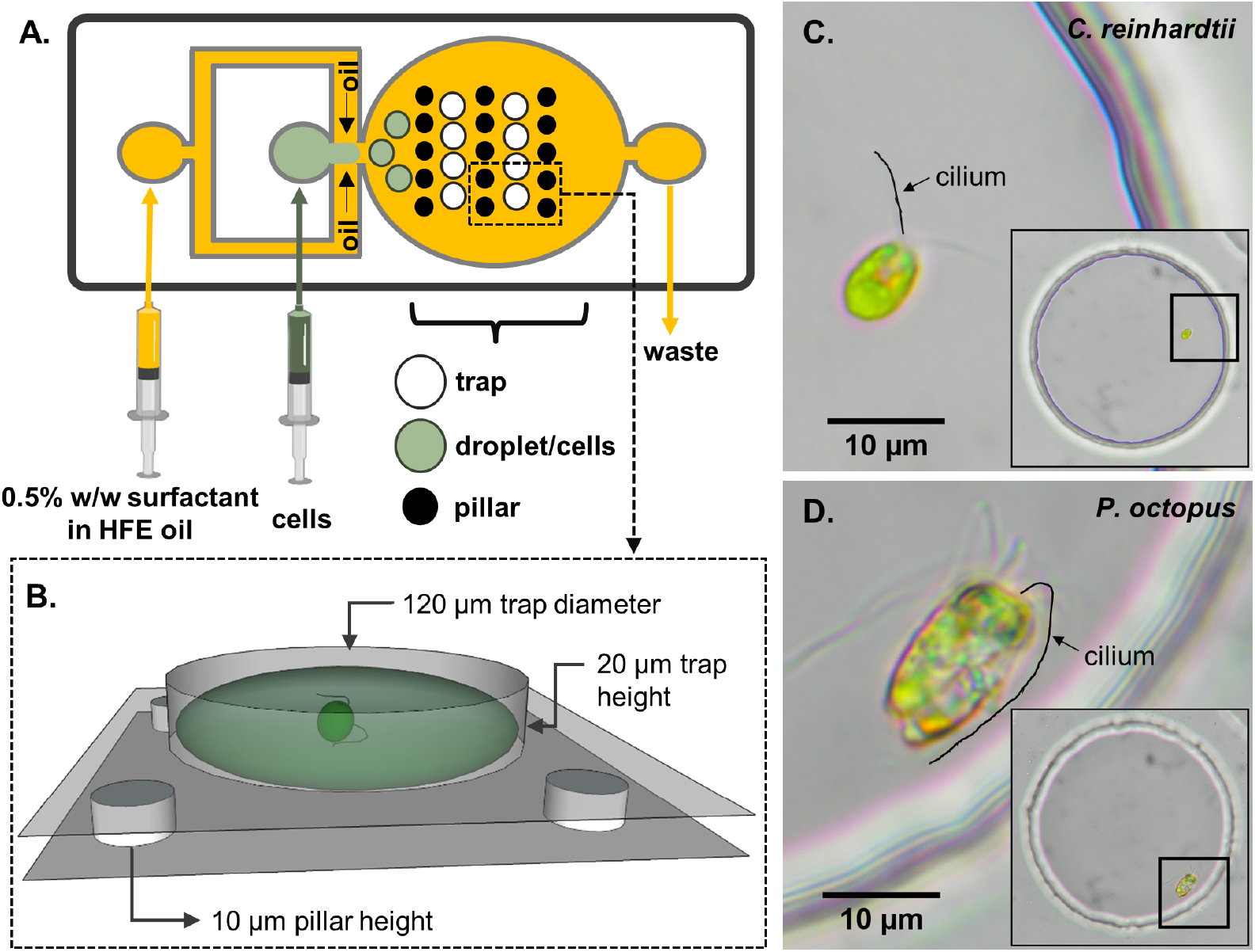
Schematic of the experimental set-up. (A) A two-layer microfluidic device with embedded single-cell traps, and syringes used for perfusion of the carrier oil phase and an aqueous suspension containing live motile cells. (B) 3D rendering of a single trap in which a cell can be stably trapped and imaged for hours. To demonstrate variability in swimming behaviour, we studied two species of motile algae, images show respectively: (C) a single *Chlamydomonas reinhardtii* (CR) cell, and (D) a single *Pyramimonas octopus* (PO) cell, in each case trapped within a 120 *μm*-diameter circular well. (Cilia positions are highlighted by manual tracings.)

The two species displayed very different behavioural signatures. Typical swimming trajectories and speeds are displayed in Figure 2. Both species have front-mounted cilia (Figure 1). CR executes a breaststroke gait where the two cilia alternate between long-periods of in-phase synchrony, and phase slips when one cilium transiently undergoes altered beating [22]. This leads to noisy swimming consisting of straight runs whenever the cilia are synchronised, and turns whenever synchrony is lost - the latter associated with reduced swimming speed [23]. PO swims forward using a rotary breaststroke with frequent episodes of ultrafast backward swimming (called ‘shocks’), where all eight cilia undulate synchronously in front of the cell for up to 20 ms [14]. With 8 beating cilia instead of 2, maximum net forward swimming speeds in PO can reach ~ 4 times that of CR (even faster during shocks: > 1 mm/s).

**Figure 2:**
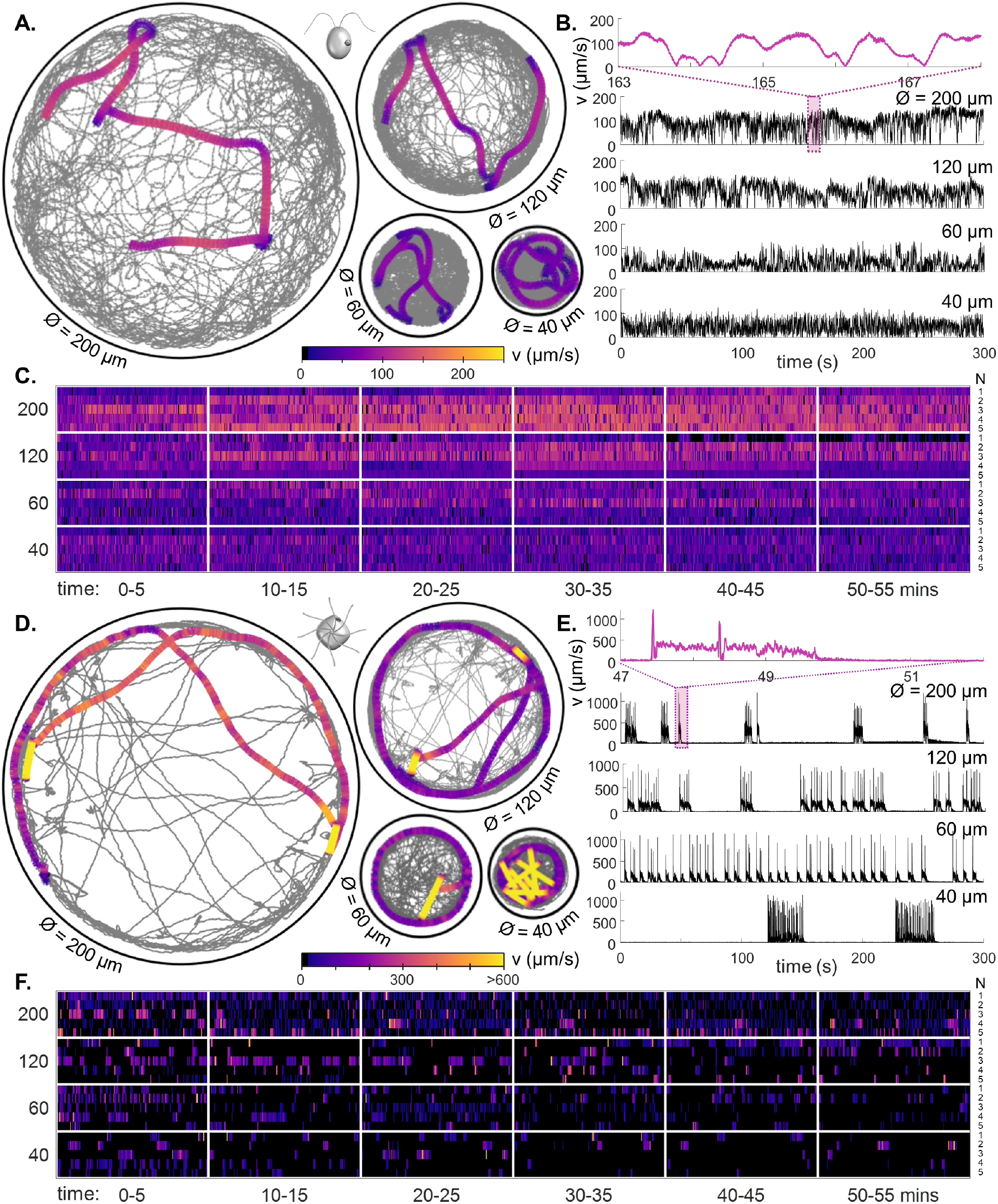
Behavioural ethograms of motile algae. (A) High-speed trajectories of single *C. reinhardtii* cells in circular traps of varying sizes (illustrating coverage over 5 minutes), overlaid with a 5-second representative trajectory (colour-coded by speed). (B) Instantaneous swimming speeds for the sample trajectories shown in (A). Inset: speed fluctuations over short timescales. (C) Heatmaps of cell swimming speed over time indexed by cell number (rows), and ordered by trap size, using data from all experimental runs. (D-F) Similar, but for PO.

The fast and episodic swimming repertoire of PO leads to a more sparse trap coverage over time (Figure 2A,D). Both species switched stochastically between different swimming modes (Figure 2B,E). While CR motility exhibits fractal characteristics (speed fluctuations at all scales), PO exhibits excitable dynamics reminiscent of neuronal spiking, where bursts of high-speed movement are followed by periods of quiescence. This distinction persists in the long-time single-cell heatmaps (Figure 2C), in which PO undergoes sharp transitions in speed and orientation compared to much smoother swimming by CR (Supplementary Videos 1, 2). In the smallest traps, PO cells showed bursting activity - with clusters of shocks in quick succession (Figure 2E). Over 1 hour, CR motility remained largely unchanged, whereas in PO a slight reduction in activity was observed in the smallest traps (Figure 2-S1).

### 2.2 Effect of physical confinement

We assayed four trap sizes, varying from strong confinement where the cell is never more than 1.5 body lengths from the boundary, to weak confinement where cells are free to roam along tight helices. Population-level trends and stereotyped behaviours were evident (Figure 2C,F) when we averaged trajectory statistics over all cells, across all timepoints.

Our findings are two-fold. First, confinement affects cell speed. As trap size is decreased, CR speeds are shifted toward lower values (Figure 3A), whereas PO decreased the characteristic speed and likelihood of runs (Figure 3B, compare location and width of the violin plot ‘waist’). With more confinement, for CR there is a marked decrease in *v* and increase in Ω, as seen in bivariate histograms of linear *v* and angular speed Ω (Figure 3C). The change in PO is more subtle: there is a reduction in the proportion of high-speed behaviour (‘run’ mode), and an increase in low-speed behaviour (‘stop’ mode) (Figure 3D). The mean-squared displacement of tracks saturates at long-times in CR to the trap radius, but not in PO. These differences indicate the radically different motility strategies of the two species (Figure 3-S1).

**Figure 3:**
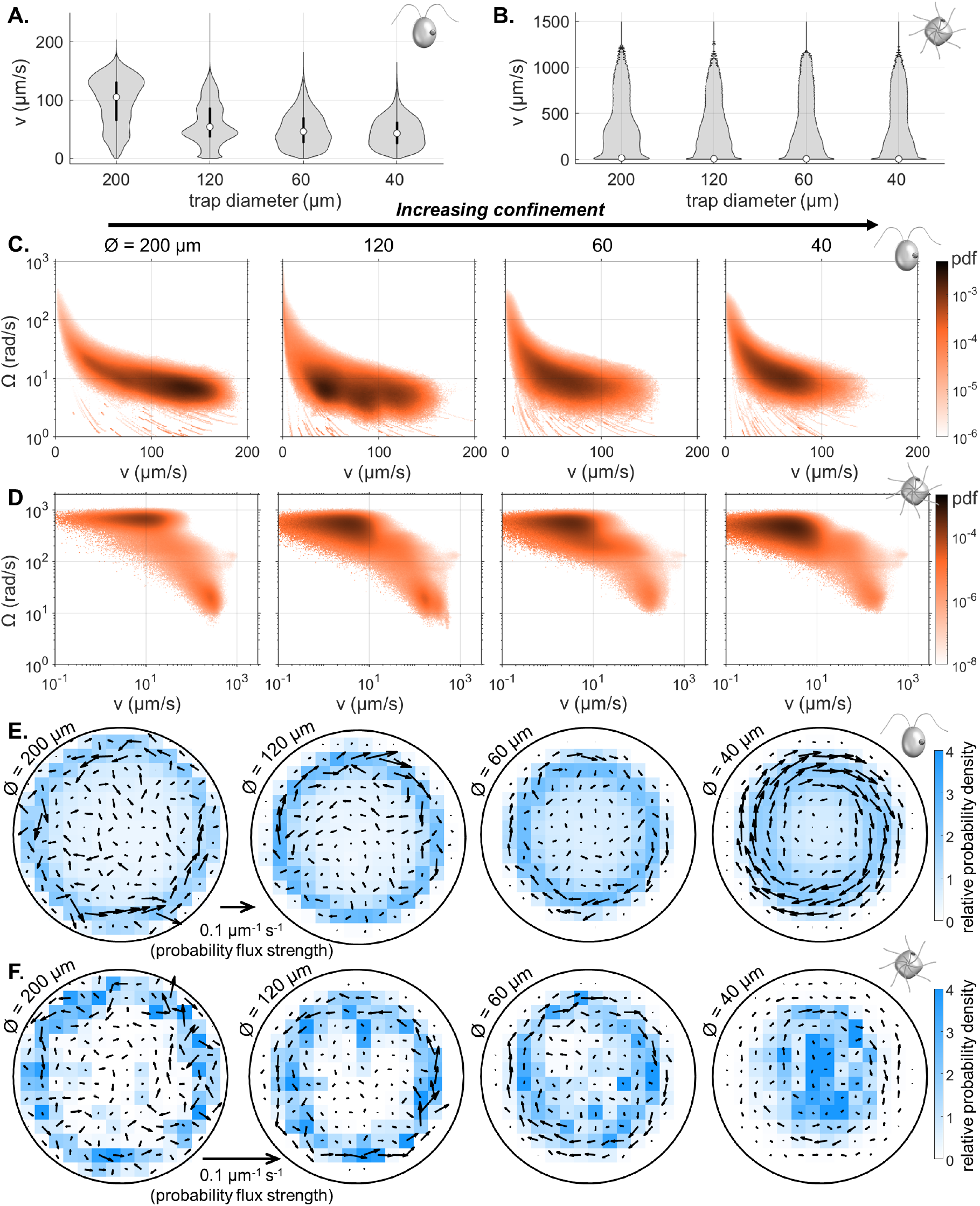
Effect of trap size on single cell motility. Violin plots of pdfs of speed (*v*) across all cells, and all time points, for CR (A) and PO (B). (Note the scaling of the pdfs is linear for CR, but logarithmic for PO.) Bivariate histograms of linear and angular speed (*v*, Ω) across all cells, and all time points for CR (C), and PO (D). Time-averaged heatmaps of the cells’ centroid positions, for each of the four different trap sizes, overlaid with arrows showing the direction and strength of steady-state non-equilibrium fluxes computed from trajectory statistics for CR (E), and PO (F).

Second, increasing confinement changes the nature of how cells explored the circular traps. To resolve these interactions in detail, we partitioned the arena into a square grid (scaled by trap size). We first computed the relative spatial probability density of occupancy in each box. Averaging over all trajectories, we found that cells were depleted from the interior of traps and instead tended to move within an annulus of the solid boundary (Figure 3E,F - shaded blue regions). This can be explained by a physical, curvature-induced effect that requires no change to the cell’s internal motility state [7].

Next, to account for the “arrow of time” in the movement history of a single individual, we computed the most probable trajectories for each trap size (see [24, 25], and Methods). We detected a chiral boundary-following behaviour that cannot be inferred from population histograms. Singlecell motility trajectories displayed steady-state non-equilibrium flux loops, becoming increasingly ordered with increasing confinement (Figure 3E,F) (particularly for CR in the smallest trap). Probability fluxes can result from asymmetries in trap curvature [25], yet here, we have discovered a novel setting in which macroscopic flux loops appear in the activity of single particles inside circular traps, i.e. without any curvature gradients (see Discussion).

### 2.3 Effect of white light stimulation

Next we explored the effect of an orthogonal parameter on motility - illumination wavelength. We illuminated the samples with broad-spectrum white light, at an intensity expected to influence photosynthesis and phototaxis in CR, but not sufficient to induced photoshock. We focused only on two trap sizes, 120 *μ*m and 60 *μ*m and again assayed over 1 hour (Supplementary Videos 3, 4).

Under white light (WL), some single-cell heterogeneity was evident (Figure 4). At early times CR swam faster in WL compared to measurements taken at in the same trap sizes at equivalent times in red light (RL), suggesting a photokinetic effect. WL also increased the likelihood of highspeed movement at early times (Figure 4A,B). A decline in motility was observed across all sampled cells after ~ 45 minutes of trapping, especially in the smallest 60 *μ*m traps (note wide ‘waists’ at zero speed). All cells in the 60 *μ*m traps were completely immotile after ~ 50 minutes.

**Figure 4:**
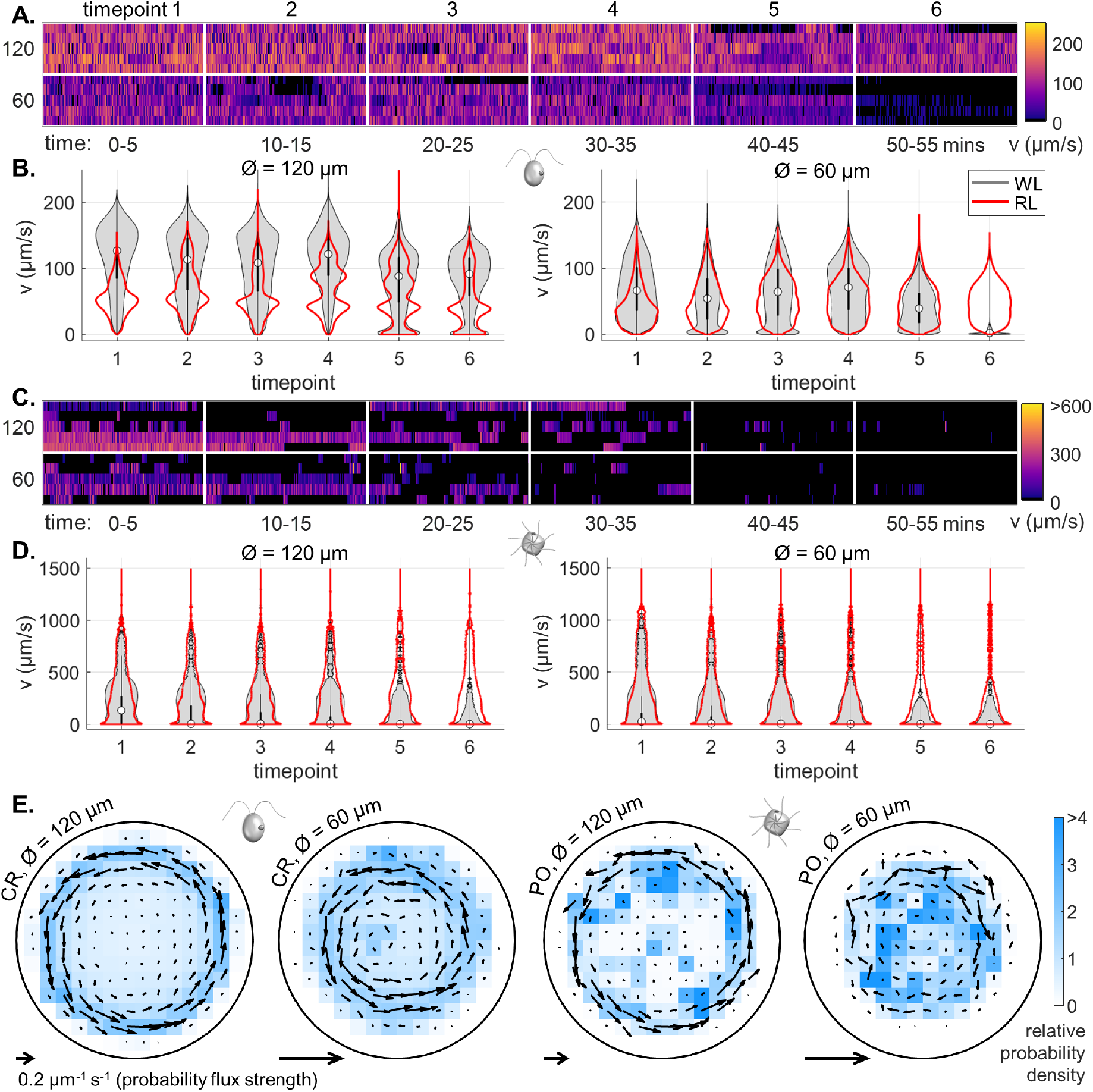
Effect of white light on single cell motility. (A) Heatmaps of cell swimming speed over time, for CR. (B) Violin plots of the pdfs (scaled linearly) of speed over time for all CR cells. (C, D) Same as (A, B) but for PO and with a logarithmic scaling for the pdfs in (D). (E) Time-averaged heatmaps of the cells’ centroid position overlaid with arrows showing the direction and strength of the probability flux computed from trajectory statistics for both trap sizes under white light conditions.

For PO, cells became increasingly quiescence and immotile over time. After one hour, all cells stopped moving (Figure 4C), and in some cases deformation of cell shape and even deciliation was noted at the final time point. By contrast in RL, only a slight reduction in motility was observed (Figure 2 and 2-S1). Comparing swimming patterns in WL and RL at early times, the maximum speeds reached (during ciliary reversals or shocks) are unchanged, but their distributions are skewed toward higher speeds due to increased likelihood of forward runs (300 – 400 *μ*m/*s*) (Figure 4D).

As before, we computed the spatial occupancy of single-cell tracks and their associated steadystate trajectory fluxes (Figure 4G,H). Directed flux loops were again revealed, most prominently in CR. For both CR and PO, the flux strengths are larger and the circulation patterns more ordered in WL at 120 *μm* than in RL at the same trap size. Interestingly for PO, the 60*μ*m traps exhibited a higher level of disorder compared to RL, likely due to lack of motility at late times.

In summary, white light increased quiescence in both species, and this effect increased with both severity of confinement *and* more time passing. The extent of motility decline under WL is therefore determined by the total duration trapped and the strength of confinement. This may reflect a cumulative, light-induced stress or phototoxicity response in green algae (see Discussion).

### 2.4 Behaviour is compressed into a trio of motility macrostates

How do we reconcile the above behavioural responses with changes in the locomotor apparatus (i.e. cilia motility)? In both species motility is inherently low-dimensional, comprising three states (Figure 5, and Appendix A): a quiescent or ‘stop’ state in which the cell body exhibits minimal movement, a ‘run’ state associated with constant speed and smooth forward swimming, and a ‘transitional’ state for (usually rapid or transient) re-orientations. Analogous motility strategies in prokaryotes include run-and-tumble in *E. coli* [26, 27], or the run-reverse-flick in *V. alginolyticus* [28]. We demonstrate how the statistics of transitions between motility macrostates can be used to provide a robust measure of behaviour that is independent of a cell’s physical environment, enabling cross-species comparisons.

**Figure 5:**
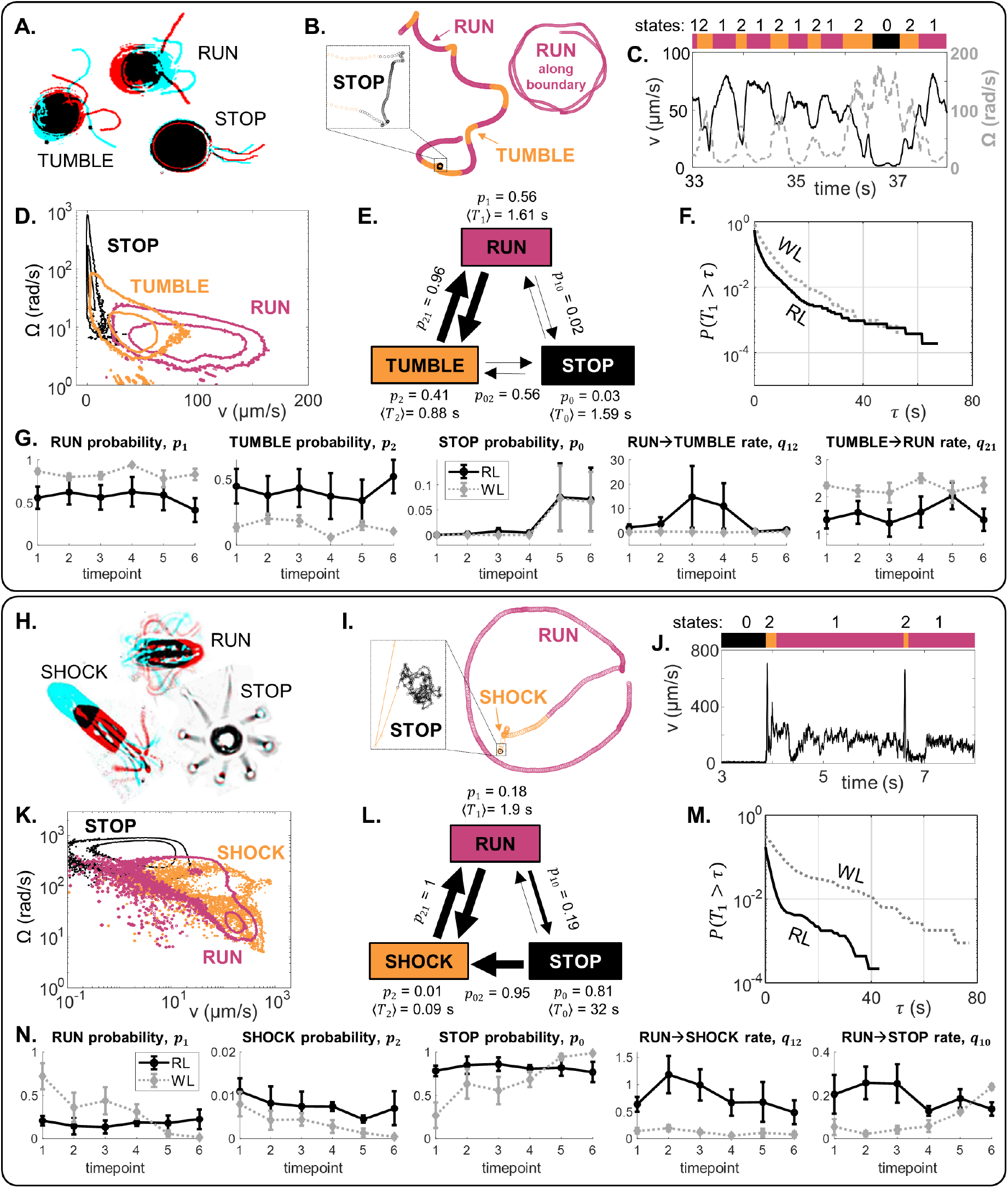
The locomotor behaviour of single-cells is controlled by a trio of motility macrostates and their transition probabilities. (A-G) For CR, distinct ciliary beat patterns and coordination states (A) produce different swimming trajectories (B), and a discrete sequence (C) of states (run,stop,tumble). For cells in 120 *μ*m traps, RL, the states occupy distinct regions in (*v*, Ω)-space (D), and a time-averaged reaction network with characteristic transition parameters (E). Network parameters are different in WL for the same trap size, as evidenced in (F): survival probability of run states, and (G): selected state probabilities and transition rates. (H-N) Similarly for PO.

Since subtle changes in motility can be hidden by population averaging, we converted individual cell tracks into discrete timeseries of three states, labelled *i* = 0, 1, 2. Swimming states are assigned according to a heuristic approach based on direct observations of ciliary beat patterns, and their associated trajectories. At any given time *t*, the cell is in state *S*(*t*). Denoting by *T_i_* the list of sojourn times in each state *i*, we estimated survival probabilities *P*(*T_i_* > *τ*), expected residence times 〈*T_i_*〉, and the pairwise state transition probabilities between states *p_ij_* (see Methods). This procedure gives rise to a unique reaction network specific to the motility strategy of each species, and each assayed condition (Figure 5E shows one such example). Rate constants are dynamic - can change in time, or in response to microhabitat perturbations. We discuss the results for each species in turn, for the 120 *μ*m traps. A similar analysis for the smaller 60 *μ*m traps can be found in Figure5-S1.

For CR, differing degrees of biciliary coordination are associated with ‘runs’, ‘slips’ or ‘drifts’ [23, 22]. We define (Figure 5A) ‘runs’ (state 1) when the two cilia engage in synchronous breaststrokes (high *v*, low Ω), ‘tumbles’ (state 2) when the cilia lose synchrony (low *v*, high Ω), and ‘stops’ (state 0) where *v* < *v_c_* for a threshold speed *v_c_* (Figure 5B,C). (‘Drifts’ are taken as generalised ‘slips’ that produce sharp turns or ‘tumbles’ due to an imbalance in the forces produced by the two cilia.) We note that while synchronous ciliary beating produces straight swimming in the trap interior (‘runs’), curvature-guided interactions with the wall lead to boundary-following behaviour (‘runs’ along the boundary); both are considered ‘runs’ here (Figure 5B).

Denoting by *p_i_* the expected probability of being in state *i*, we found that in the 120 *μ*m traps (Figure 5F), 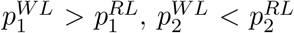 (Figure 5F). These differences in CR motility WL or RL in the 120 *μ*m traps are also stable over time (Figure 5G). In contrast 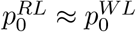 (Figure 5G). WL increased the tumble→run transition rate (*q*_21_) and decreased the run → tumble transition rate (*q*_12_) (Figure 5G), where *q_ij_* denotes the transition rate from state *i* to *j*. Empirically, this means that CR performs more frequent and longer runs in WL compared to RL.

The motility of PO can also be mapped to a tripartite run-stop-shock repertoire [14], where PO ‘shocks’ are analogous with CR ‘tumbles’. ‘Runs’ result from coordinated breaststrokes involving 8-cilia [6], meanwhile ‘shocks’ occur when all eight cilia switch to a symmetric beat and undulate in front of cell, producing rapid backward swimming followed by reorientation (Figure 5H). As in CR, both forward swimming across the chamber and circling at the boundary are classified as runs (Figure 5I). Here, ‘runs’ (state 1) correspond to moderate *v* and low Ω, ‘shocks’ (state 2) to high *v* and high *d*Ω/*dt*, and ‘stops’ (state 0) are where *v* < *v_c_* for a threshold speed *v_c_* (Figure 5J,K).

Again, we computed rate constants to determine the structure of the PO motility network, e.g. Figure 5L. Comparing PO behaviour in WL and RL for the same trap size (Figure 5M), we found that state probabilities *p*_0,1,2_ remained largely constant over 1 hour in RL, but changed over time in WL. Initially, the probability of observing a run state is higher in WL, but after the 4th timepoint (about 40 minutes) this drops to below the corresponding RL value. We found that 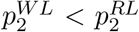, and 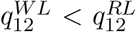. In WL, both *p*_0_, *q*_10_ increased over time. This means that in WL, PO runs are longer and more likely than in RL, but cells lost motility over time (more likely to stop).

### 2.5 Algal cell motility is light-switchable

Since light induced behavioural changes in both species, we next asked how the *same* individual responds to photostimulation, and if such responses are reversible. The following dynamic assay was performed. Single CR or PO cells were trapped in 120 *μ*m wells and imaged continuously for 20 minutes in the following sequence: 1) 5 minutes of RL (RL1), 2) remove red filter, image for 5 minutes (WL1), 3) replace red filter, image for a further 10 minutes (RL2). For both species, motility in RL1 and WL1 was similar to previous results for constant RL or WL illumination. Most cells swam faster in each prospective WL1 period than in the preceding RL1 period (Figure 6A,B).

Both species displayed a reversible but asymmetric photokinetic response to 5 minutes of WL-stimulation. Response to RL → WL (step-up) is fast and occurs within seconds, but recovery after WL → RL (step-down) is slow and persists for several minutes into the RL2 phase (Figure 6C,D). CR in particular, produced a stronger speed response to step-up stimulation than to the opposite. For PO, the increase in mean speed is mainly due to increased probability runs. In WL1, the likelihood of state transitions in both species decreased (Figure 6D), as in continuous WL. This is particularly noticeable in PO, which responds immediately to step-up by almost completely suppressing state transitions. Cells gradually *adapted* by increasing the transition frequency over the remainder of WL1, and well into RL2. Thus, algal swimming motility is light-switchable in the presence of short-lived stimuli, which induce perturbations to the cell’s internal motility state network. Cells retained a *memory* of the stimulation (WL1) for longer than the stimulus duration itself.

**Figure 6:**
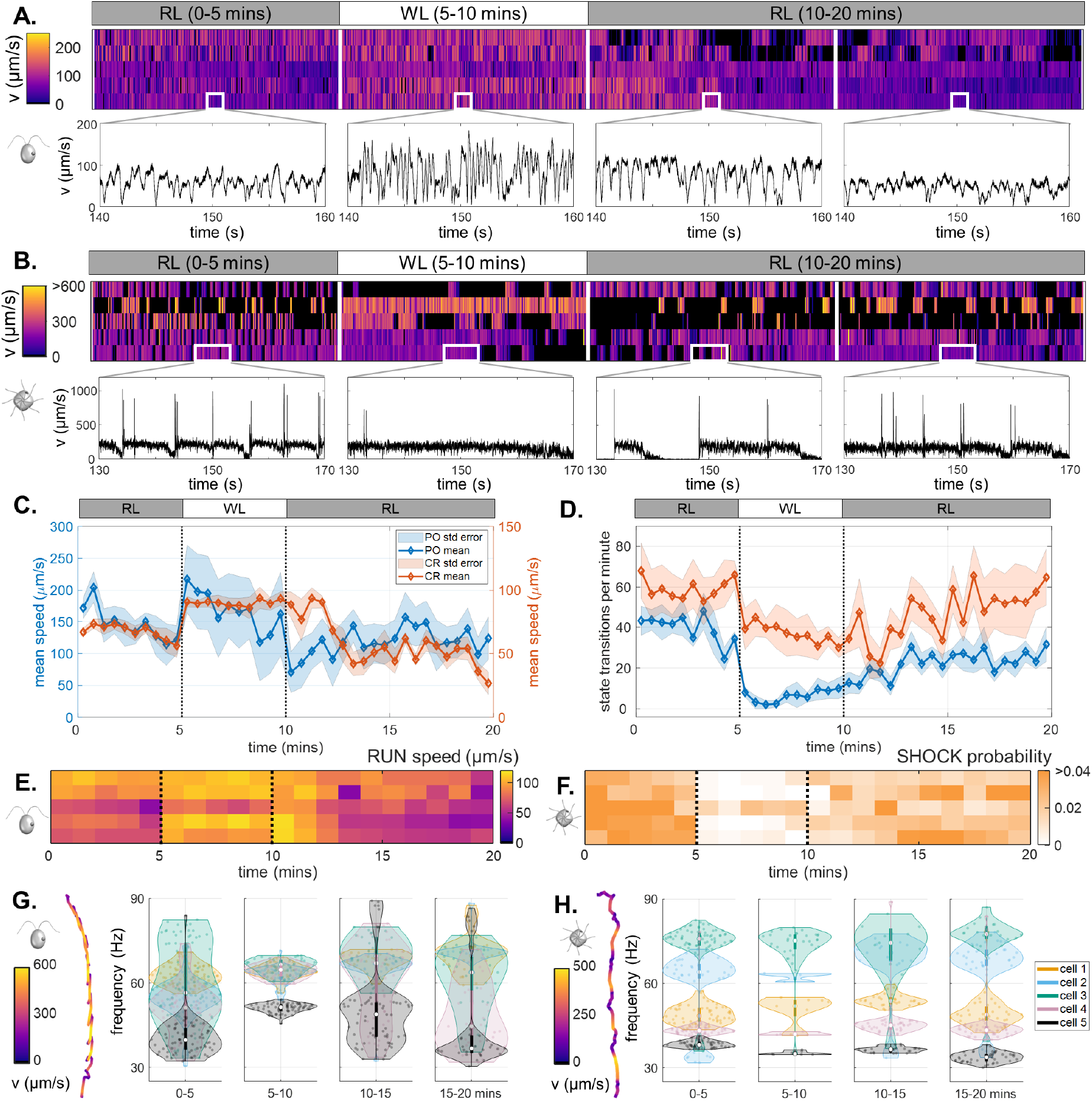
Light-switchable motility of green algae. (A) Heatmaps of CR swimming speed over the course of a 20-minute photoresponse assay, insets show short, representative timeseries in each phase. (B) The same for PO. (C) Population-averaged swimming speed, over time. (D) Population-averaged rates of transitions between motility states, over time. (Shaded regions are standard errors.) (E) Mean speed of ‘runs’ in CR, over time. (F) Probability of ‘shocks’ in PO, over time. Violin plots of the most prominent frequencies extracted from helical trajectories (inset), for CR (G) and for PO (H). For CR, this corresponds to the ciliary beat frequency.

In WL1, CR increased the speed of prospective runs (Figure 6E), but PO suppressed the likelihood of shocks (Figure 6F). Analysing ‘run’ phases exclusively, we further deduced that CR swims faster by increasing the frequency *and* synchronicity of ciliary beating (from 52 ± 4 to 62 ± 3 Hz), in response to step-up from RL1 to WL1 (Figure 6G). A similar analysis of PO trajectories yielded significant cell-to-cell variability but no clear frequency signatures that could be correlated with ciliary beating (Figure 6H). This is likely due to real-time changes in coordination of the 8 cilia leading to asymmetries in propulsive force that can alter the helicity of trajectories [29]. This highlights a notable difference between CR and PO photoresponses to WL-stimulation: while CR changes swimming speed, PO changes state transitions.

### 2.6 Rapid chemical modification of motility triggered by droplet fusion

In many protists, ciliary motility is modulated by chemical sigalling, including Ca^2+^-influx through ion channels, suggesting a conserved Ca^2+^-dependent mechanism [30, 31]. Ionic fluxes elicit transitions in cilia beating, waveform and swimming behaviour [32, 33, 34, 35]. In *Paramecium*, Ca^2+^, and K^+^ ions are major regulators of motility, the latter inducing episodes of backward swimming [36]. We hypothesized that motile algae also possess a behavioural response to K^+^, and confirmed this in PO with a bulk motility assay (see Methods). The introduction of a droplet pre-loaded with 10 mM KCl to a suspension of PO cells produced perturbed swimming (Figure 7A).

**Figure 7:**
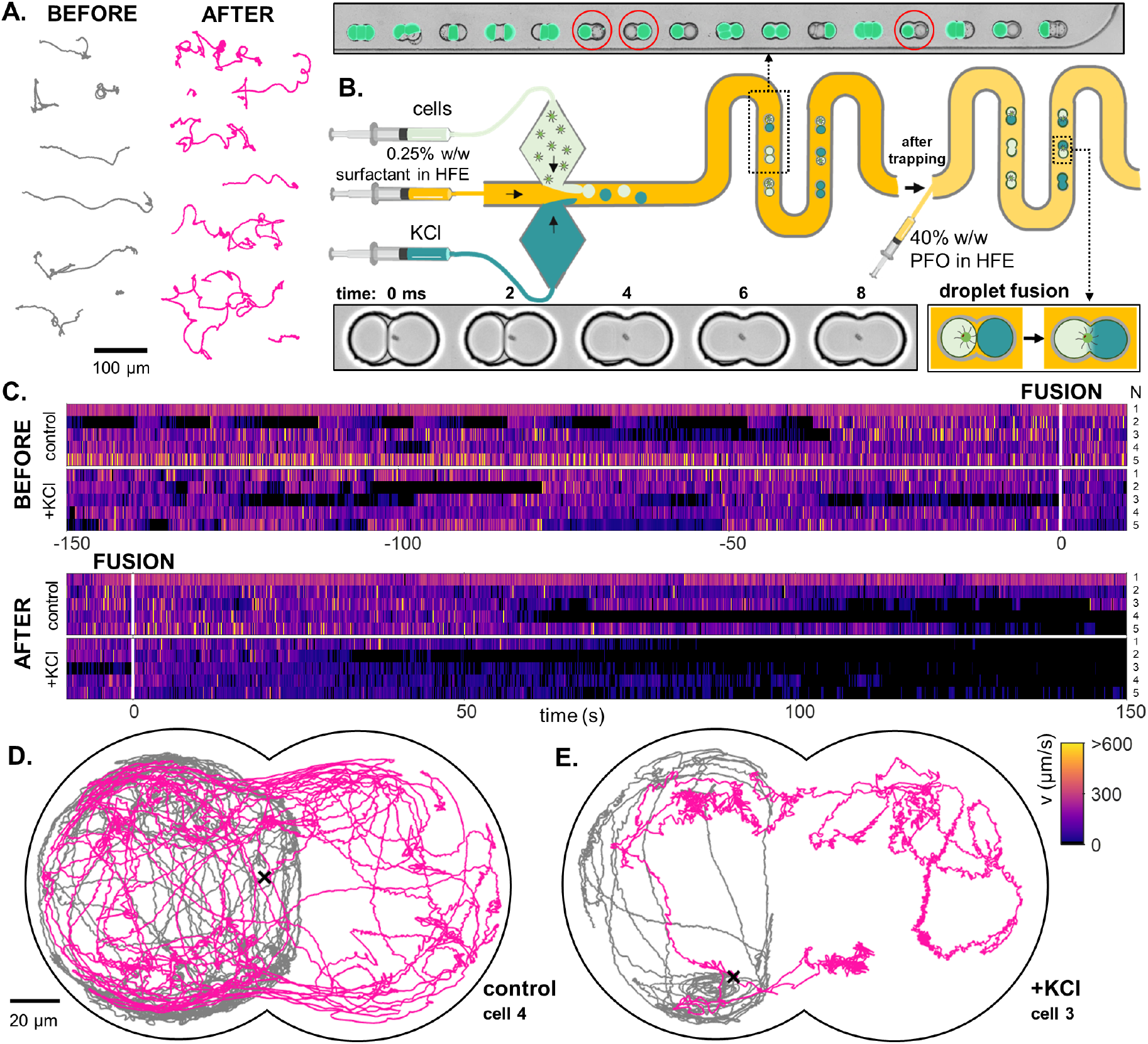
A novel droplet-fusion assay for querying single-cell motility responses to rapid chemical perturbation. (A) Samples tracks from a bulk motility assay (see Methods), immediately *before* (‘normal motility phenotype’) and *after* addition of 10 mM KCl to the medium (‘high KCl phenotype’). (B) Schematic of the serpentine microfluidic device for generation and pairing of droplets. Fluorescein was added to the KCl phase to identify paired droplets - one containing the trapped cell, and the other loaded with KCl (inset, red circles). Fusion was then induced by replacing the 0.5% surfactant in HFE with 40% 1H,1H,2H,2H-Perfluoro-1-octanol (PFO) in HFE. Two such droplets merge within 1 or 2 ms (inset). (C) Heatmaps of single-cell swimming speed, before and after the moment of fusion to a droplet containing KCl, compared to a control pair where KCl was replaced with inert culture medium. An example single-cell trajectory before (grey) and after (pink) droplet fusion, for a control (D) and a KCl pair (E).

However, bulk assays do not allow path-sampling of single-cell behaviour. Instead we designed a microfluidic device to deliver on-demand fusion of paired droplets, allowing us to study for the first time how cell motility changes in response to *instantaneous* chemical perturbations (Figure 7B). Contrary to traditional assays that use flow channels to deliver concentration gradients, our design applies real-time control and localised perturbations to single microbes. Using a cross-junction microfluidic chip, we generated in alternation droplets containing either KCl or cell suspension. Pairs of droplets were then trapped in downstream doublet wells. We then identified droplet pairs comprising one droplet with the trapped PO cell, and another droplet containing the assay medium supplemented with 10 mM KCl and 1 *μ*M of fluorescein which acted as a reporter. Fusion was triggered by starting the flow of 40% perfluoro-1-octanol (PFO) in HFE (a competing surfactant) (see Methods). Two such droplets fused within 1 or 2 ms. Note that not all droplet pairs generated will have the target composition, since trap-occupancy was not perfectly made up of alternating droplets (Figure 7B, inset).

We compared the motility of cells in these doublet traps in the presence or absence of KCl. Control experiments were performed by substituting the chemical assay medium with normal culture medium (without cells). Prior to fusion, cell motility was similar in both cases (Figure 7C). After fusion, cells in the KCl droplets immediately showed erratic swimming, followed by gradual decay in motility over time (see Supplementary Video 5). Significant changes in the morphology and tortuosity of swimming trajectories was also observed after fusion (Figure 7D,E). Further examples of this response may be found in Figure 7-S1.

## 3 Discussion

### 3.1 Stereotypy and the ‘arrow of time’ in long-term single-cell behaviour

The emerging field of computational ethology seeks to track and annotate the complex natural behaviours of organisms [37, 38, 39, 40]. Due to their small size, variable morphology, and high speed movement, detailed analysis of the behavioural patterns of microorganisms has been technically challenging. Here we leveraged droplet microfluidics to track the swimming trajectories of microbes for long periods of time. We compared a freshwater biflagellate with a marine octoflagellate, and evaluated single-individual responses to a large number of controlled environmental stimuli (including mechanical, light, chemical).

Our results showed that the behavioural space of roaming microbes comprises only a small set of stereotyped movements. This inherent low-dimensionality was observed in other organisms [41, 42, 43] and artificial microswimmers [44, 45]. We discovered a minimal set of three motility macrostates: a canonical ‘run’ state, a ‘stop’ or rest state, and a transitory state (‘tumbles’ or ‘shocks’) involving trajectory reorientations, and used this to propose a new paradigm for phenotyping microbial motility. Microorganisms inhabiting different ecological niches likely evolved divergent strategies for sensorimotor integration, despite conserved cilia-based motility. For example, CR swims according to a more continuous repertoire, whereas PO motility was episodic or excitable, fluctuating stochastically between fast swimming and behavioural quiescence, reminiscent of transitions between ‘wakeful’ and ‘sleep states’ observed in many animals [46]. Future work should seek to map the shape space of permissible ciliary waveforms [35] and any multiciliary actuation patterns [6], to the resulting species-specific motility behaviours.

Importantly, our lifetime behavioural recordings retained the ‘arrow of time’ [47]. This gives us access to key information that cannot otherwise be derived from traditional, bulk-averaged measurements, including chiral movements in confined spaces (see next section). From these timeseries, we reconstructed internal motility networks for single cells in terms of their motility state-transition probabilities, showing how the latter changes in time but also in response to environmental stimuli. Ultralong timeseries can also be used to infer non-equilibrium entropy production [48] and irreversibility [14] in the motility repertoire of single cells. We hypothesize that these distinctive, species-specific movement signatures reflect intracellular signalling processes controlling the algal ciliary motility apparatus [49], and could be key to the emergence of cognition and environmental responsiveness in protists [50, 31].

### 3.2 Non-equilibrium flux loops without curvature gradients

Microswimmers often interact with their physical environment, particularly with interfaces and boundaries. Microscale cell-environment interactions are commonplace in natural settings with exposed surface features or other heterogeneous structures, for example soil, foam, or particulate matter [9, 8, 51]. Here, we engineered PDMS chambers with precise shapes and geometries to explore how confinement affects cell motility. In red light, cells tolerated long-periods of confinement with little change in their overall motility characteristics. Both CR and PO cells exhibited a ‘circling’ movement along boundaries, consistent with a previous study suggesting that this arises from interactions between puller-type swimming (forward ‘runs’) near a solid boundary with curvature [7]. Circling is a direct consequence of physical confinement and does not require any change of internal swimming state.

In the smallest traps and under white light illumination, we also discovered a novel chiral movement (Figures 3 & 4). While confinement and edge effects has been shown previously to induce or stabilise macroscopic chiral movement in swimmer suspensions [52, 53], here chirality has emerged in the travel history of a *single cell*. This provides a distinct route towards a chiral nonequilibrium steady-state that has not been reported previously, since CR-like swimmers roaming inside circular traps with constant curvature were suggested to be incapable of producing flux loops on average [25]. This discrepancy highlights the importance of further particulars of cell shape and/or swimming mechanism [29, 54, 55, 56]. Further experimental and modelling efforts are currently underway to resolve these microswimmer-boundary interactions [57], including the role of cilia mechanosensitivity [14].

### 3.3 Light-dependent algal motility and phototoxicity

Comparing the algae’s long-term motility we found that swimming in RL is more stable than in WL. Short periods of WL exposure increases the average run-speed of CR by increasing ciliary beat frequency, consistent with previous observations on micropipette-immobilised cells [58]). In contrast, WL decreases the frequency of gait transitions in PO, producing longer stops, long runs, and fewer shocks. Prolonged or excessive exposure (>40 minutes) to WL eventually led to reduced or cessation of motility in both species. In CR this is likely due to WL-induced adhesion [59]. In PO, some cells showed an irreversible structural disintegration suggestive of photodamage - occasionally individual cilia can detach from the cell body. Increasing confinement (smaller traps) decreased the time taken to reach this state, suggesting that motility deterioration may be due to buildup of some metabolite or excretion from the trapped cell over time [60].

Shorter periods (~ 5 minutes) of WL-stimulation produced reversible changes in motility in the algae (Figure 6), again with species-dependent photokinetic responses occurring at the single cell level. While CR responded largely by modulating run speed, PO modulated the balance of motility macrostates (e.g. suppressed shocks). These distinct behavioural responses offer intriguing prospects for synthetic biology and bioengineering, such as light-guided patterning of photoresponsive microbial [61, 62]. Going forward, it will be interesting to combine our assay with genetic perturbations (particularly in CR), or design light intensity gradients within a trap, to reveal the regulatory pathways responsible for photosynthesis, phototransduction, and phototaxis.

### 3.4 Future prospects of droplet microfluidics for assaying cell motility

Micro-encapsulation technologies provide unique functionalities that enable controlled, on-demand creation and manipulation of mimic micro-environments [28]. A microfluidics-based pipeline allowed us to stably trap motile cells, and to establish their unique behavioural signatures. We use multilayer devices and static droplet trapping wells [19] to keep droplets stable for hours, and isolated from possible fluctuations in the continuous phase [63]. This allowed us to achieve long-time imaging, and reliable control over trap size, shape and geometry. The trapping arrays we used also maximised the probability of obtaining multiple single-trapped cells in a single experiment, thus increasing analytical throughput. Our current design has focused on imaging a small number of traps at a time, simultaneous, high-throughput imaging of multiple cells in multiple wells is also possible, but at the detriment of spatial resolution and thus may not be sufficient for identifying motility state or behavioural transitions.

Our work paves the way for new applications and opportunities for designing diagnostic tools in the absence of molecular tests. Microfluidics-based assays enable the detection of single-cell motility signatures and any heterogeneity in microbial behaviour. Phenotypic diversity is critical in microbial ecosystems, where they may give rise to distinct selection pressures and antimicrobial resistance [64]. Our droplet fusion device represents a highly-novel solution for assaying fast cell-environment interactions and responses to chemical perturbations (Figure 7). Further developments and extensions are on the way, including augmentation of on-demand single cell encapsulation with active cell sorting [65, 66]. The integration of lab-on-chip technologies, high-speed microscopy and computer vision has significant potential for reconstructing the species-specific sensorimotor pathways of microorganisms, and revealing their response thresholds to dynamic environmental perturbations.

## 4 Materials and Methods

### 4.1 Cell culturing and maintenance

*C. reinhardtii* cultures were prepared from axenic plates of the CC125 WT strain (Chlamydomonas Center). Individual colonies were transferred to 25 cm^3^ volumes of Tris-minimal media. Liquid cultures were grown under a 14/10 light-dark cycle at 21°C and 40% humidity, with constant shaking at 110 rpm. Liquid cultures were sub-cultured when they approached the end of the exponential phase. For motility experiments, second- or third-generation cultures were harvested in late-exponential phase (6-9 days after inoculation). Cell density was measured as 1 × 10^6^ cells per mL. Cells were centrifuged at 100g for 10 minutes, and then concentrated 10-fold. The cells were then left in darkness for a minimum of 30 minutes to dark-adapt them before each experimental run.

We prepared our PO cultures from axenic liquid cultures of the WT of the species *P. octopus* (NIVA/NORCCA). 200 μL of axenic culture was transferred to 25 cm^3^ of TL30 media. Cells were grown under continuous illumination at 21 °C and 40% humidity, without shaking. For motility experiments, cultures were harvested during the latter half of the exponential phase (20 - 30 days after inoculation). Cell density was measured at 3 × 10^3^ cells per ml. Cells were centrifuged at 100g for 10 minutes, and then concentrated 10-fold. The cells were then left in darkness for a minimum of 30 minutes to dark-adapt them before each experimental run.

### 4.2 Microfluidic chip fabrication

The devices were designed with CAD software (DraftSight, Dassault Systems) and fabricated following classical soft-lithography procedures by using a high-resolution acetate mask (Microlithography Services Ltd.). Negative photoresist SU-8 3025 (MicroChem, Newton, MA) was spin-coated onto clean silicon wafers to a thickness of 10 *μ*m, patterned by exposure to UV light through the photomask (Xia and Whitesides, 1998) and hard bake at 95 °C for ~ 7 minutes.. Prior to development through immersion in propylene glycol monomethyl ether acetate (PGMEA, Sigma-Aldrich), a second layer of SU-8 3025 at 20 *μ*m in height was spin-coated, UV exposed and hard baked (95 °C, ~ 7 minutes) for the development of the trapping arrays. Uncured polydimethylsiloxane (PDMS) consisting of a 10:1 polymer to cross-linker mixture (Sylgard 184) was poured onto the master, degassed, and baked at 70 °C for 4 hours. The PDMS mould was then cut and peeled from the master, punched with a 1.5 mm biopsy punch (Kai Medical) to create inlet ports for tubing insertion. A total of three holes were punched; two inlets for the continuous and aqueous phase and an outlet for waste collection. For the time-lapse device, the PDMS mould was plasma bonded to thin cover slips (22 × 50 mm, 0.13 – 0.17 mm thick). Hydrophobic surface treatment was performed immediately after bonding by flushing with 1% (v/v) Trichloro (1H, 1H, 2H, 2H-perfluorooctyl) silane (Aldrich) in HFE-7500, and placed in a 65 °C oven for 30 min.

### 4.3 Flow-focusing droplet generation

Microfluidic device fabrication was done using classical soft lithography techniques. A total of 4 devices were developed. All devices consisted of a flow-focusing junction for droplet generation (1A) (height, 10 *μm*), and a second layer with a trapping array made up of circular wells (height, 20 *μm*). The trapping sizes developed were 40, 60, 120 and 200 *μ*m. The dimensions of the flow-focusing junction varied depending on trap size. We designed a range of dimensions for the flow focusing junctions and matching trap sizes (i.e. circle diameter). Depending on the trapping size, the total number of traps was between 78 (∅ 200 *μm*) to 840 (∅ 40 *μm*).

Droplets were typically generated at rates approx ~ 50 per second. The flow rates were controlled using syringe pumps (Nemesys, Cetoni), 1 mL plastic syringes (BD PlastipakTM; sterile needles, 25G x 1” – NR. 18, 0.5 mm x 25 mm, BD MicrolanceTM 3), and portex tubing PE (Scientific Laboratory Supplies, 0.38 × 0.355 mm). The flow rates for oil and cell suspension were varied depending the size dimensions of each device. A 1:3 ratio was aimed for the continuous and aqueous phase, respectively.

The carrier oil phase was prepared using fluorinated oil HFE-7500 (Fluorochem Ltd) containing 0.5% (w/v) 008-Fluorosurfactant (RAN Biotechnologies, Inc.). The aqueous phase consisted of liquid cell cultures (see Cell culturing and maintenance). The droplets were generated at the flow-focusing junction creating water-in-oil emulsions. A dilute suspension of algae was injected through the inlet. Following trapping, droplets were stably confined to the microwell during imaging acquisition. Once all the traps were filled, the aqueous phase flow was halted and the continuous phase was flowing at a reduced (5x) rate to flush away excess droplets.

### 4.4 Live-cell high-speed imaging

Brightfield imaging was conducted with an inverted microscope (Leica Microsystems, DMi8), equipped with a high-speed camera (Phantom Vision Research, V1212). We first scanned the array of trapped cells to locate traps matching our criteria (droplet fitting exactly into the trap, droplet containing only one cell). For the 40 μm, 60 μm and 120 μm trap sizes, as well as the cell fusion experiments, we used a 20x long-working distance objective (HC PL/0.40). For the largest 200 *μ*m traps, we lowered the magnification to 5x (NPLAN/0.12) equipped with a 1.6x tube lens, to reduce file size. All traps were imaged with the same intensity and aperture settings, and at 500fps. For the 1 hour confinement experiments, cells were imaged continuously but 5 minute recordings taken at 5 minute intervals, to obtain a total of 6 timepoints per cell. Data from droplets that were disrupted at any point during imaging was discarded.

### 4.5 Light-modulation experiments

For brightfield imaging in WL, we used a standard broad-spectrum LED source to illuminate the specimen. Red light (RL) imaging was accomplished by insertion of an IR long-pass filter (610nm, Chroma) to the light path. Spectra corresponding to the two possible illumination options are compared in Figure8. For light-switching experiments, the red filter was removed or inserted manually.

**Figure 8:**
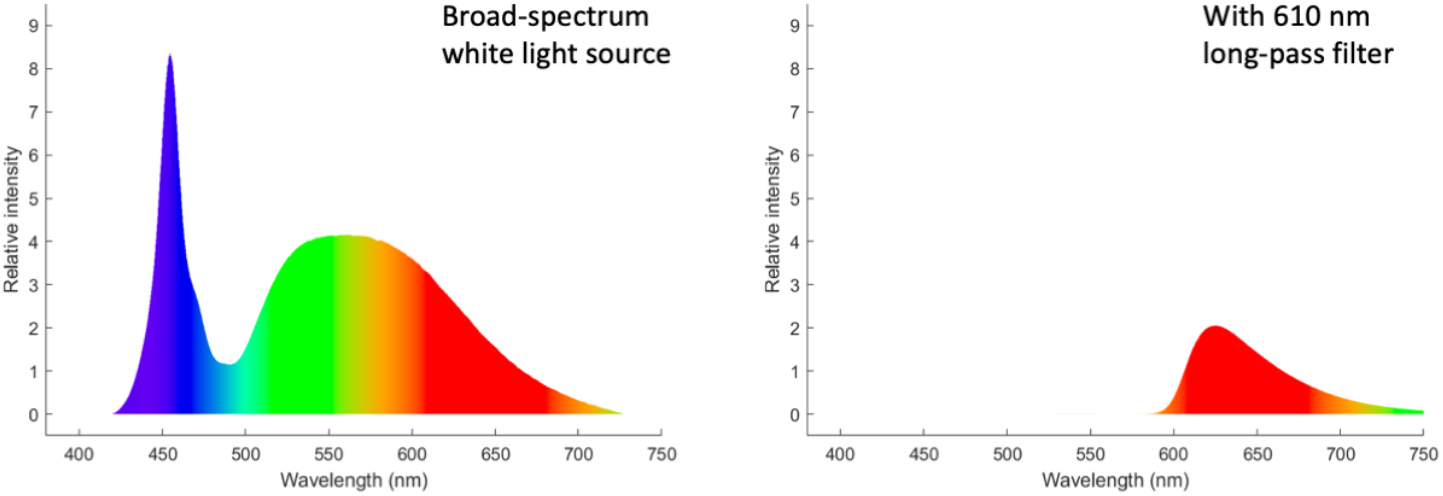
Illumination spectra recorded with a spectrometer (OceanInsight OceanHDX, 200*μ*m-fibre).

### 4.6 Bulk cell motility assay

We assayed the effect of KCl on *P. octopus* behaviour via a simple open-air method. We first added 1 μL of a concentrated suspension of cells to a glass coverslip under red light in a dark room. We waited 5s for flows from the placement of the droplet to subside, before imaging for 25s (at 10x, 100fps). We then added either a 1 μL droplet of culture media (for the control) or a 1 μL drop of 50mM KCl (for the KCl test), and after waiting another 5s for flows to subside we imaged for another 25s.

### 4.7 Paired-droplet fusion assay

Droplet pairs were generated using a cross junction microfluidic device in which cells and KCl solutions flowing in separate channels are encapsulated in alternation. We used a 25 degrees angle previously reported to produce the most stable alternation function. When stable alternation was achieved, droplets were suddenly halted by removal of the inlet tubings of KCl and cell solutions followed by gentle removal of excess droplets at 3 *μ*l/min for ~ 1 minute. We subsequently identified droplet pairs of expected volumes with one containing a single cell. To ensure fusion with KCl and the absence of mixing prior to fusion, the 10 mM KCl solution was spiked with 1 *μ*M fluorescein (Merck) which was imaged before triggering of fusion. A 15 minute video of the cells was acquired displaying the entrapped droplets 7.5 minutes prior and 7.5 minutes post fusion. Fusion was induced by surfactant replacement with 1H,1H,2H,2H-Perfluoro-1-octanol (Merck) (PFO). A solution of 40% PFO in HFE was flown at 5 *μ*l/min and run until fusion was achieved (~ 4 min). PFO competes with the fluorosurfactant which destabilizes the droplet interface to induce rapid, reproducible fusion. Control experiments without KCl were done to confirm the absence of confounding factors.

For the fusion experiments, the identification process consisted of two steps. Firstly we located a trap that had two equal droplets in place, one containing a cell. Secondly, we took a fluorescence image to verify that the droplet without the cell contained KCl. The fluorescence image was taken in the LASX software, using a broad-spectrum LED source (CoolLED-pE300) equipped with a triple-band filter set (including FITC, Ex: 475 nm, Em: 530 nm). The fluorescence intensity was set to 60%, the exposure time was 600 ms, and the gain was 2.0. The presence of fluorescence covering the whole of the empty droplet was sufficient to prove the presence of KCl.

**Table 1:** A comparison of the concentrations of key ions (in mM), before and after paired droplet fusion.

| Ion | Pre-fusion | Post-fusion (with KCl) |
| --- | --- | --- |
| Na<sup>+</sup> | 136.3 | 68.2 |
| K<sup>+</sup> | 3.0 | 6.5 |
| Ca<sup>2+</sup> | 2.9 | 1.5 |
| Mg<sup>2+</sup> | 13.8 | 6.9 |
| Cl<sup>-</sup> | 161.5 | 80.8 |
| SO<sub>4</sub><sup>2-</sup> | 4.8 | 2.4 |
| NO<sub>3</sub><sup>-</sup> | 1.1 | 0.6 |
| HPO<sub>4</sub><sup>2-</sup> | 0.1 | negligible |

### 4.8 Image processing and cell tracking

Raw video data was exported to 8-bit grayscale and enhanced by subtracting an average image in MATLAB (Mathworks). Trap boundaries were identified manually to increase the fidelity of 2D cell tracking, which was performed automatically using the Trackmate plugin in ImageJ [67]. A Laplace of Gaussian detector was used for spot identification, with slightly different blob diameters for CR and for PO (14 um and 21 um respectively). Single continuous tracks were obtained for each experimental run (N=5 individuals per condition), and exported for further processing and extraction of detailed track features/other statistics (see Appendix A). Video frames from the bulk motility assays were processed and analysed similarly.

## 5 Author contributions

SAB, VA performed experiments and acquired data; HLS, SAB, KYW analysed and modelled data. KYW, FG designed experiments, acquired funding, and supervised the study. KYW conceived the project, and wrote the original draft of the paper. All authors were involved in reviewing and editing.

## 6 Acknowledgments

This work received funding from the European Research Council (ERC) under the European Union’s Horizon 2020 research and innovation programme (grant agreement No 853560 *EvoMotion*, to KYW), and a Springboard Award from the Academy of Medical Sciences and Global Challenges Research Fund (*SBF*003\1160 to KYW). The research was also funded by the Biological and Biotechnological Research Council (grant BB/T011777/1 to FG).

## 7 Resource availability

The following datasets were generated [link upon publication]. All codes associated with this manuscript can be found [link upon publication].

## References

[1] H. S. Jennings, “On the Significance of the Spiral Swimming of Organisms,” The American Naturalist, vol. 35, pp. 369–378, May 1901.

[2] G. J. Berman, “Measuring behavior across scales,” BMC Biology, vol. 16, p. 23, Dec. 2018.

[3] A. Mathis, P. Mamidanna, K. M. Cury, T. Abe, V. N. Murthy, M. W. Mathis, and M. Bethge, “DeepLabCut: markerless pose estimation of user-defined body parts with deep learning,” Nature Neuroscience, vol. 21, pp. 1281–1289, Sept. 2018.

[4] A. E. X. Brown, E. I. Yemini, L. J. Grundy, T. Jucikas, and W. R. Schafer, “A dictionary of behavioral motifs reveals clusters of genes affecting Caenorhabditis elegans locomotion,” Proceedings of the National Academy of Sciences, vol. 110, pp. 791–796, Jan. 2013.

[5] S. M. Coyle, E. M. Flaum, H. Li, D. Krishnamurthy, and M. Prakash, “Active systems encode an emergent hunting behavior in the unicellular predator lacrymaria olor.,” Current biology : CB, vol. 29, pp. 3838–3850.e3, Nov. 2019.

[6] K. Y. Wan, “Synchrony and symmetry-breaking in active flagellar coordination,” Philosophical Transactions of the Royal Society B-biological Sciences, vol. 375, p. 20190393, Feb. 2020.

[7] T. Ostapenko, F. J. Schwarzendahl, T. J. Böddeker, C. T. Kreis, J. Cammann, M. G. Mazza, and O. Bäumchen, “Curvature-Guided Motility of Microalgae in Geometric Confinement,” Physical Review Letters, vol. 120, p. 068002, Feb. 2018.

[8] V. Kantsler, J. Dunkel, M. Polin, and R. E. Goldstein, “Ciliary contact interactions dominate surface scattering of swimming eukaryotes,” Proceedings of the National Academy of Sciences, vol. 110, pp. 1187–1192, Jan. 2013.

[9] A. Théry, Y. Wang, M. Dvoriashyna, C. Eloy, F. Elias, and E. Lauga, “Rebound and scattering of motile *Chlamydomonas* algae in confined chambers,” Soft Matter, p. 10.1039.D0SM02207A, 2021.

[10] T. Bhattacharjee and S. S. Datta, “Bacterial hopping and trapping in porous media,” Nature Communications, vol. 10, p. 2075, Dec. 2019.

[11] T. A. G. Hageman, M. P. Pichel, P. A. Löthman, J. Cho, M. Choi, N. Korkmaz, A. Manz, and L. Abelmann, “Long-term observation of Magnetospirillum gryphiswaldense in a microfluidic channel,” Archives of Microbiology, vol. 201, pp. 1427–1433, Dec. 2019.

[12] A. Codutti, M. A. Charsooghi, E. Cerdá-Doñate, H. M. Taïeb, T. Robinson, D. Faivre, and S. Klumpp, “Single-cell motion of magnetotactic bacteria in microfluidic confinement: interplay between surface interaction and magnetic torque,” preprint, Biophysics, Mar. 2021.

[13] S. Sasso, H. Stibor, M. Mittag, and A. R. Grossman, “From molecular manipulation of domesticated Chlamydomonas reinhardtii to survival in nature,” eLife, vol. 7, p. e39233, Nov. 2018.

[14] K. Y. Wan and R. E. Goldstein, “Time Irreversibility and Criticality in the Motility of a Flagellate Microorganism,” Physical Review Letters, vol. 121, p. 058103, Aug. 2018.

[15] D. B. Brückner, A. Fink, C. Schreiber, P. J. F. Röttgermann, J. O. Räadler, and C. P. Broedersz, “Stochastic nonlinear dynamics of confined cell migration in two-state systems,” Nature Physics, vol. 15, pp. 595–601, June 2019.

[16] L. Tweedy, P. A. Thomason, P. I. Paschke, K. Martin, L. M. Machesky, M. Zagnoni, and R. H. Insall, “Seeing around corners: Cells solve mazes and respond at a distance using attractant breakdown,” Science, vol. 369, p. eaay9792, Aug. 2020.

[17] X. Niu, F. Gielen, A. J. deMello, and J. B. Edel, “Electro-Coalescence of Digitally Controlled Droplets,” Analytical Chemistry, vol. 81, pp. 7321–7325, Sept. 2009.

[18] L. Frenz, J. Blouwolff, A. D. Griffiths, and J.-C. Baret, “Microfluidic Production of Droplet Pairs,” Langmuir, vol. 24, pp. 12073–12076, Oct. 2008.

[19] E. Fradet, C. McDougall, P. Abbyad, R. Dangla, D. McGloin, and C. N. Baroud, “Combining rails and anchors with laser forcing for selective manipulation within 2d droplet arrays,” Lab Chip, vol. 11, pp. 4228–4234, 2011.

[20] H. Harz and P. A. Hegemann, “Rhodopsin-regulated calcium currents in Chlamydomonas,” Nature, vol. 351, pp. 489–491, 1991.

[21] G. Kreimer, “The green algal eyespot apparatus: a primordial visual system and more?,” Curr Genet, p. 25, 2009.

[22] K. Y. Wan, K. C. Leptos, and R. E. Goldstein, “Lag, lock, sync, slip: the many ‘phases’ of coupled flagella,” Journal of The Royal Society Interface, vol. 11, p. 20131160, May 2014.

[23] M. Polin, I. Tuval, K. Drescher, J. P. Gollub, and R. E. Goldstein, “Chlamydomonas Swims with Two “Gears” in a Eukaryotic Version of Run-and-Tumble Locomotion,” Science, vol. 325, pp. 487–490, July 2009.

[24] C. Battle, Broedersz Chase P., Fakhri Nikta, Geyer Veikko F., Howard Jonathon, Schmidt Christoph F., and MacKintosh Fred C., “Broken detailed balance at mesoscopic scales in active biological systems,” Science, vol. 352, pp. 604–607, Apr. 2016.

[25] J. Cammann, F. J. Schwarzendahl, T. Ostapenko, D. Lavrentovich, O. Bäumchen, and M. G. Mazza, “Emergent probability fluxes in confined microbial navigation,” Proceedings of the National Academy of Sciences, vol. 118, p. e2024752118, Sept. 2021.

[26] H. C. Berg, “The Rotary Motor of Bacterial Flagella,” Annual Review of Biochemistry, vol. 72, pp. 19–54, June 2003.

[27] E. Perez Ipiña, S. Otte, R. Pontier-Bres, D. Czerucka, and F. Peruani, “Bacteria display optimal transport near surfaces,” Nature Physics, vol. 15, pp. 610–615, June 2019.

[28] K. Son, D. R. Brumley, and R. Stocker, “Live from under the lens: exploring microbial motility with dynamic imaging and microfluidics,” Nature Reviews Microbiology, vol. 13, pp. 761–775, Dec. 2015.

[29] D. Cortese and K. Y. Wan, “Control of Helical Navigation by Three-Dimensional Flagellar Beating,” Physical Review Letters, vol. 126, p. 088003, Feb. 2021.

[30] K. Inaba, “Calcium sensors of ciliary outer arm dynein: functions and phylogenetic considerations for eukaryotic evolution.,” Cilia, vol. 4, p. 6, 2015.

[31] K. Y. Wan and G. Jékely, “Origins of eukaryotic excitability,” Philosophical Transactions of the Royal Society B-biological Sciences, vol. 376, p. 20190758, 2021.

[32] C. Beck and R. Uhl, “On the localization of voltage-sensitive calcium channels in the flagella of Chlamydomonas reinhardtii.,” The Journal of cell biology, vol. 125, pp. 1119–1125, June 1994.

[33] C. Kung and Y. Naito, “Calcium-induced ciliary reversal in the extracted models of “Pawn”, a behavioral mutant of Paramecium.,” Science (New York, N.Y.), vol. 179, pp. 195–196, Jan. 1973.

[34] R. Brette, “Integrative Neuroscience of Paramecium, a “Swimming Neuron”.,” eNeuro, vol. 8, June 2021.

[35] V. F. Geyer, J. Howard, and P. Sartori, “Ciliary beating patterns map onto a low-dimensional behavioral space that accords with a simple mechanochemical model,” arXiv:2106.00704 [cond-mat], 2021.

[36] I. Kunita, S. Kuroda, K. Ohki, and T. Nakagaki, “Attempts to retreat from a dead-ended long capillary by backward swimming in Paramecium,” Frontiers in Microbiology, vol. 5, p. 270, 2014.

[37] S. Han, E. Taralova, C. Dupre, and R. Yuste, “Comprehensive machine learning analysis of Hydra behavior reveals a stable basal behavioral repertoire,” eLife, vol. 7, p. e32605, Mar. 2018.

[38] O. Yaski, J. Portugali, and D. Eilam, “Arena geometry and path shape: When rats travel in straight or in circuitous paths?,” Behavioural Brain Research, vol. 225, pp. 449–454, Dec. 2011.

[39] E. Yemini, T. Jucikas, L. J. Grundy, A. E. X. Brown, and W. R. Schafer, “A database of Caenorhabditis elegans behavioral phenotypes,” Nature Methods, vol. 10, pp. 877–879, Sept. 2013.

[40] G. J. Stephens, M. Bueno de Mesquita, W. S. Ryu, and W. Bialek, “Emergence of long timescales and stereotyped behaviors in Caenorhabditis elegans,” Proceedings of the National Academy of Sciences, vol. 108, pp. 7286–7289, May 2011.

[41] B. T. Larson, J. Garbus, J. B. Pollack, and W. F. Marshall, “A unicellular walker embodies a finite state machine,” Biorxiv, 2021.

[42] D. Jordan, S. Kuehn, E. Katifori, and S. Leibler, “Behavioral diversity in microbes and lowdimensional phenotypic spaces,” Proceedings of the National Academy of Sciences, vol. 110, pp. 14018–14023, Aug. 2013.

[43] A. C. H. Tsang, A. T. Lam, and I. H. Riedel-Kruse, “Polygonal motion and adaptable phototaxis via flagellar beat switching in the microswimmer Euglena gracilis,” Nature Physics, vol. 14, pp. 1216–1222, Dec. 2018.

[44] M. Leoni, M. Paoluzzi, S. Eldeen, A. Estrada, L. Nguyen, M. Alexandrescu, K. Sherb, and W. W. Ahmed, “Surfing and crawling macroscopic active particles under strong confinement: Inertial dynamics,” Physical Review Research, vol. 2, p. 043299, Dec. 2020.

[45] B. V. Hokmabad, R. Dey, M. Jalaal, D. Mohanty, M. Almukambetova, K. A. Baldwin, D. Lohse, and C. C. Maass, “Emergence of Bimodal Motility in Active Droplets,” Physical Review X, vol. 11, p. 011043, Mar. 2021.

[46] D. E. Lawler, Y. L. Chew, J. D. Hawk, A. Aljobeh, W. R. Schafer, and D. R. Albrecht, “Sleep Analysis in Adult C. elegans Reveals State-Dependent Alteration of Neural and Behavioral Responses,” Journal of Neuroscience, vol. 41, no. 9, pp. 1892–1907, 2021.

[47] F. S. Gnesotto, F. Mura, J. Gladrow, and C. P. Broedersz, “Broken detailed balance and nonequilibrium dynamics in living systems: a review,” Reports on Progress in Physics, vol. 81, p. 066601, June 2018.

[48] D. J. Skinner and J. Dunkel, “Improved bounds on entropy production in living systems,” Proceedings of the National Academy of Sciences, vol. 118, p. e2024300118, May 2021.

[49] H. Guo, Y. Man, K. Y. Wan, and E. Kanso, “Intracellular coupling modulates biflagellar synchrony,” Journal of The Royal Society Interface, vol. 18, p. 20200660, Jan. 2021.

[50] I. Kunita, T. Yamaguchi, A. Tero, M. Akiyama, S. Kuroda, and T. Nakagaki, “A ciliate memorizes the geometry of a swimming arena,” Journal of The Royal Society Interface, vol. 13, p. 20160155, May 2016.

[51] M. Souzy, A. Allard, J.-F. Louf, M. Contino, I. Tuval, and M. Polin, “Microbial narrow-escape is facilitated by wall interactions,” 2021.

[52] H. Wioland, F. G. Woodhouse, J. Dunkel, J. O. Kessler, and R. E. Goldstein, “Confinement Stabilizes a Bacterial Suspension into a Spiral Vortex,” Physical Review Letters, vol. 110, p. 268102, June 2013.

[53] K. Beppu, Z. Izri, T. Sato, Y. Yamanishi, Y. Sumino, and Y. T. Maeda, “Edge current and pairing order transition in chiral bacterial vortices,” Proceedings of the National Academy of Sciences, vol. 118, p. e2107461118, Sept. 2021.

[54] E. Lauga, W. R. DiLuzio, G. M. Whitesides, and H. A. Stone, “Swimming in Circles: Motion of Bacteria near Solid Boundaries,” Biophysical Journal, vol. 90, pp. 400–412, Jan. 2006.

[55] E. Lushi, V. Kantsler, and R. E. Goldstein, “Scattering of biflagellate microswimmers from surfaces,” Physical Review E, vol. 96, p. 023102, Aug. 2017.

[56] M. Zaferani, F. Javi, A. Mokhtare, and A. Abbaspourrad, “Effect of flagellar beating pattern on sperm rheotaxis and boundary-dependent navigation,” preprint, Biophysics, Jan. 2020.

[57] S. E. Spagnolie, C. Wahl, J. Lukasik, and J.-L. Thiffeault, “Microorganism billiards,” Physica D: Nonlinear Phenomena, vol. 341, pp. 33–44, Feb. 2017.

[58] U. Rüffer and W. Nultsch, “Flagellar photoresponses of *Chlamydomonas* cells held on micropipettes: I. Change in flagellar beat frequency: Flagellar Beat Frequency Changes,” Cell Motility and the Cytoskeleton, vol. 15, no. 3, pp. 162–167, 1990.

[59] C. T. Kreis, M. Le Blay, C. Linne, M. M. Makowski, and O. Bäumchen, “Adhesion of Chlamydomonas microalgae to surfaces is switchable by light,” Nature Physics, vol. 14, pp. 45–49, Jan. 2018.

[60] J. Boedicker, M. Vincent, and R. Ismagilov, “Microfluidic Confinement of Single Cells of Bacteria in Small Volumes Initiates High-Density Behavior of Quorum Sensing and Growth and Reveals Its Variability,” Angewandte Chemie, vol. 121, pp. 6022–6025, June 2009.

[61] G. Frangipane, D. Dell’Arciprete, S. Petracchini, C. Maggi, F. Saglimbeni, S. Bianchi, G. Vizsnyiczai, M. L. Bernardini, and R. Di Leonardo, “Dynamic density shaping of photokinetic E. coli.,” eLife, vol. 7, Aug. 2018.

[62] M. R. Bittermann, D. Bonn, S. Woutersen, and A. Deblais, “Light-switchable deposits from evaporating drops containing motile microalgae,” arXiv:2105.00270 [cond-mat], May 2021. arXiv: 2105.00270.

[63] S. H. Han, J. Kim, and D. Lee, “Static array of droplets and on-demand recovery for biological assays,” Biomicrofluidics, vol. 14, p. 051302, Sept. 2020.

[64] N. Dhar and J. D. McKinney, “Microbial phenotypic heterogeneity and antibiotic tolerance.,” Current opinion in microbiology, vol. 10, pp. 30–38, Feb. 2007.

[65] V. Anagnostidis, B. Sherlock, J. Metz, P. Mair, F. Hollfelder, and F. Gielen, “Deep learning guided image-based droplet sorting for on-demand selection and analysis of single cells and 3D cell cultures,” Lab on a Chip, vol. 20, no. 5, pp. 889–900, 2020.

[66] L. Howell, V. Anagnostidis, and F. Gielen, “Multi-object detector yolov4-tiny enables high-throughput combinatorial and spatially-resolved sorting of cells in microdroplets,” Advanced Materials Technologies, 2021.

[67] J.-Y. Tinevez, N. Perry, J. Schindelin, G. M. Hoopes, G. D. Reynolds, E. Laplantine, S. Y. Bednarek, S. L. Shorte, and K. W. Eliceiri, “TrackMate: An open and extensible platform for single-particle tracking.,” Methods (San Diego, Calif.), vol. 115, pp. 80–90, Feb. 2017.

[68] B. Bechtold, P. Fletcher, seamusholden, and S. Gorur-Shandilya, “bastibe/violinplot-matlab: A good starting point,” Feb. 2021.

[69] N. Tarantino, J.-Y. Tinevez, E. F. Crowell, B. Boisson, R. Henriques, M. Mhlanga, F. Agou, A. Israël, and E. Laplantine, “TNF and IL-1 exhibit distinct ubiquitin requirements for inducing NEMO-IKK supramolecular structures,” Journal of Cell Biology, vol. 204, pp. 231–245, 01 2014.

