## Appendix for "Phenotyping single-cell motility in microfluidic confinement"

### 8 Appendix A: Analysis of trajectories

Trajectories obtained from cell tracking (see Materials and Methods) consist of 2D coordinates at discrete times ($t_1$, $t_2$, $t_3$...) with a constant time interval, i.e. $\Delta t = t_{i+1} - t_i = 0.002s$ for videos captured at 500 fps. The trajectories were analysed using custom MATLAB scripts. First, a smoothing filter was applied. For CR, a Savitzky–Golay filter with order 2 and frame length 201 was used to smooth out the helical trajectories such that a speed and angular velocity corresponding to the net forward motion of the cell could be obtained. For PO, a Savitzky–Golay filter with order 2 and frame length 21 was used. A smaller frame length was used due to the fast timescale of the shock behaviour and since PO has smoother trajectories than CR (Appendix 1 Figure 9). The cell velocity at time $t_i$ was calculated as

$$\mathbf{v}(t_i) = \frac{\mathbf{x}(t_{i+1}) - \mathbf{x}(t_i)}{\Delta t}, \tag{1}$$

where $\mathbf{x}(t_i)$ is the 2D coordinate of the centroid of the cell at time $t_i$ for the smoothed trajectories. The angular velocity of the cell was defined as

$$\Omega(t_i) = \frac{\arccos \hat{\mathbf{v}}(t_{i-1}) \cdot \hat{\mathbf{v}}(t_i)}{\Delta t}, \tag{2}$$

where $\hat{\mathbf{v}}(t_i)$ is the normalised velocity vector at time $t_i$. To reduce the noise of the angular velocity data, the results reported are a moving mean across 25 frames.

Violin plots, which combine box plot and histogram data representations into one diagram, were created using a MATLAB package [67]. The width of the violin plots correspond to the probability density function and are scaled linearly for CR and logarithmically for PO.

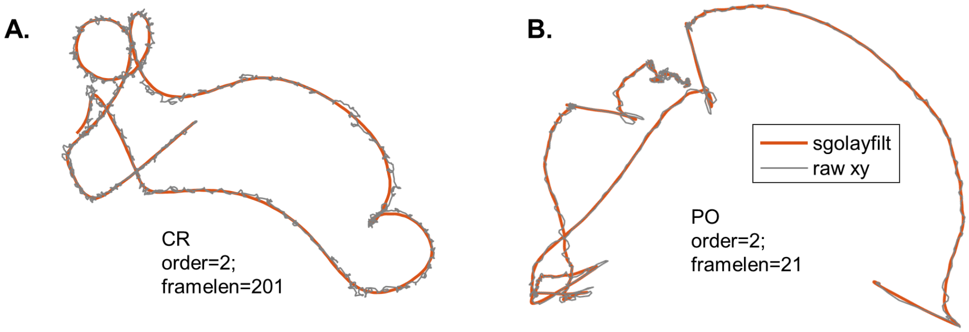


Figure 9: Smoothing tracking data using a Savitzky-Golay filter (sgolayfilt). Example raw and smoothed trajectories for CR (A) and PO (B).

#### 8.1 Mean square displacement

The the mean square displacement (MSD) is defined as

$$\mathrm{MSD}(\tau) = \frac{1}{N_\tau} \sum_{t_i} |\mathbf{x}(t_i + \tau) - \mathbf{x}(t_i)|^2, \tag{3}$$

where $\tau$ is the delay and $N_\tau$ the number of pairs of time points in the trajectory with $\Delta t = \tau$. The MSD was calculated for the raw 2D coordinate positions using the '@msdanalyzer' MATLAB package [68]. The MSD results reported here were calculated using 1/50th of the data points (i.e. every 50th frame) due to the data array size constraints of the package.

#### 8.2 Probability flux calculation and relative probability density

We calculated the probability fluxes using a method previously applied to trajectories of *C. reinhardtii* [24] and first introduced by [23]. The 2D positional space is divided into a grid of equally sized square boxes with side length $\Delta x = r_{trap}/7$. For each time point the position is assigned to a box $(i,j)$ , where $i$ and $j$ are the box positions in the $x$ and $y$ direction respectively. From these coarse-grained trajectories, a time-series of transitions is obtained by constructing the following array

$$A = \begin{bmatrix} (i,j)_1 & (i,j)_2 & t_{1,2} \\ (i,j)_2 & (i,j)_3 & t_{2,3} \\ ... & ... & ... \\ (i,j)_{N-1} & (i,j)_N & t_{N-1,N} \end{bmatrix}, \tag{4}$$

where $(i,j)_n$ and $(i,j)_{n+1}$ are the positions of consecutively visited boxes and $t_{n,n+1}$ is the length of time spent in the initial state $(i,j)_n$ before transitioning to the new state $(i,j)_{n+1}$. In a small number of cases, the two success states $(i,j)_n$ and $(i,j)_{n+1}$ do not correspond to nearest neighbours. In such cases, the intermediate boxes are determined by linear interpolation and the corresponding extra transitions are inserted into the array $A$ (equation 4) to ensure that all transitions in $A$ are between nearest neighbours.

The net transition rates between neighbouring boxes are calculated from the coarse-grained trajectories using

$$\omega_{(i,j)(k,l)} = \frac{1}{t_{total}} \left( N_{(i,j)(k,l)} - N_{(k,l)(i,j)} \right), \tag{5}$$

where $N_{(i,j)(k,l)}$ is the number of transitions from box $(i,j)$ to box $(k,l)$ and $t_{total}$ is the total duration of the trajectory. The net transition rates are then used to calculate the probability flux

$$\mathbf{j}_{(i,j)} = \frac{1}{2\Delta x} \begin{pmatrix} \omega_{(i,j)(i+1,j)} + \omega_{(i-1,j)(i,j)} \\ \omega_{(i,j)(i,j+1)} + \omega_{(i,j-1)(i,j)} \end{pmatrix}. \tag{6}$$

This method is summarised in Appendix 1 Figure 10. Note that this calculation does not account for diagonal transitions, however since they account for only $< 5\%$ of all nearest neighbour transitions, we assume they have a minimal effect on the results.

The 2D trajectory data was also used to calculate the relative probability density

$$c_{(i,j)} = \frac{A_{trap} n_{(i,j)}}{A_{box} \sum n_{(i,j)}}, \tag{7}$$

where $n_{(i,j)}$ is the number of trajectory points within box $(i,j)$, $A_{box} = \Delta x^2$ is the box area and $A_{trap} = \pi r_{trap}^2$ is the area of the trap [6].

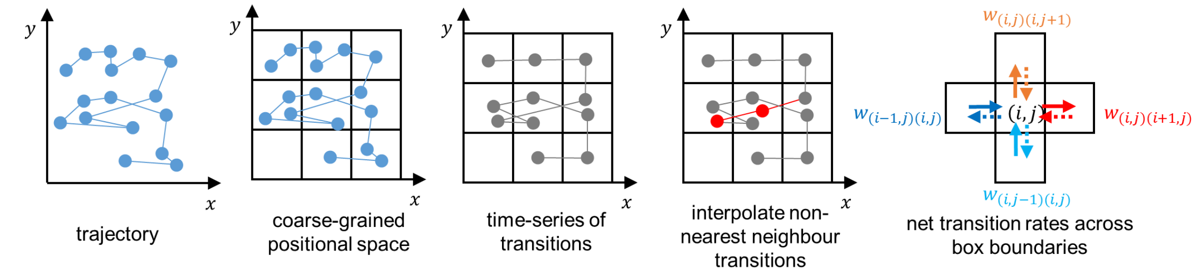


Figure 10: Probability flux analysis method.

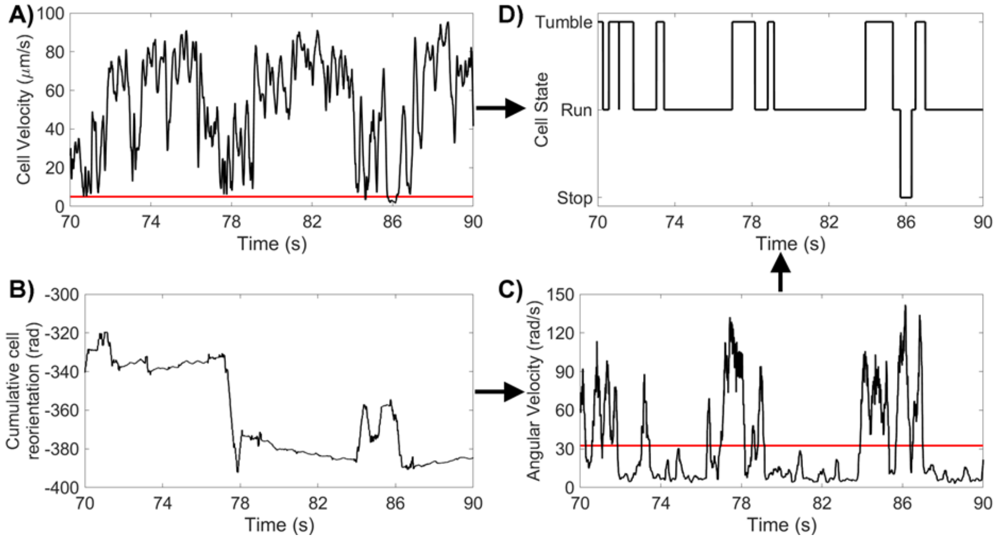


Figure 11: Method For Identifying CR States

A) CR cell velocity for a 10 s segment, with a run/stop threshold of 5 µm/s. B) CR cumulative reorientation for the same 10 s segment. C) CR angular velocity for the same 10 s segment, with a tumble threshold of 32.5 rad/s. D) CR states for the same 10 s segment.

### 9 Conversion into three motility macrostates

We started from raw track data, and assigned states in CR and PO using a combination of linear $v$ and/or angular $\Omega$ speeds. In both cases, we took moving averages to reduce frame-to-frame noise (due to the high imaging frame rates).

#### 9.1 States for *Chlamydomonas reinhardtii*: (run, stop, tumble)

We defined the 'stop' state as times when $v <$5 µm/s. A smoothing filter was applied to the binary stop/move data to remove spurious state transitions. We then verified the stop states by visual inspection. To identify 'tumbles', we use angular velocity to locate times when $\Omega >$32.5 rad/s. We filtered the dateset to correct for false tumbles by removing tumbles that were under 200ms, and for false runs of very short duration found between successive tumbles. We again verified that tumble

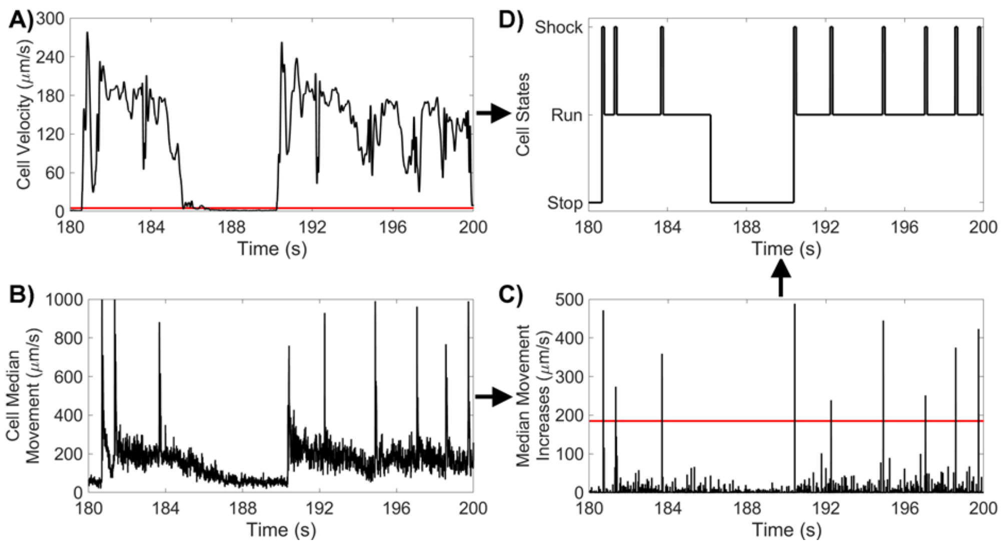


Figure 12: Method For Identifying CR States
A) PO cell velocity for a 10 s segment, with a run/stop threshold of 5 µm s$^{-}$1. B) PO median movement over 9-frame windows for the same 10 s segment. C) PO increases in cell median movement for the same 10 s segment, with a shock threshold of 185 µm/s. D) PO states for the same 10 s segment.

states corresponded with cell reorientation with visual inspection. Finally, frames that were neither a ‘stop’ or a ‘tumble’, were designated ‘runs’, with ‘stops’ taking precedence over ‘tumbles’. The workflow is summarised in Figure11.

#### 9.2 States for *Pyramimonas octopus*: (run, stop, shock)

Here, only the linear speed is sufficient to assign states to PO, since they are associated with very distinct speeds [13]. We defined the ‘stop’ state as times when the smoothed speed $v$ <5 µm/s. A smoothing filter was applied to the binary stop/move data to remove spurious state transitions. We then verified the stop states by visual inspection.

To identify shocks, we first smoothed the data by computing a local median value and then a moving mean. We then identified each local minimum and the subsequent local maximum, and computed the increase between the two. Increases of $\Omega$ >185 µm/s were identified as shocks. The start point of shocks were chosen as the point in time between the local minimum and maximum where half of the total increase in displacement had occurred. Equivalently, the end of the shock was defined as the point in time between the current displacement maximum and post-shock minimum where half of the total decrease in displacement had occurred. Finally, frames that were neither a ‘stop’ or a ‘shock’, were designated ’runs’, with ‘stops’ taking precedence over ‘shocks’. Brief (¡0.2s) ‘run’ states between a ‘stop’ and a ‘shock’ were reclassified as ‘stop’ states to remove spurious state transitions. The workflow is summarised in Figure 12.

##### 9.3 State probability and transition rate analysis

We estimated transition probabilities between motility macrostates via a simple counting algorithm. State probability is given by

$$p_i = \frac{n_i}{\sum_j n_j}, \tag{8}$$

where $n_i$ is the number of frames in which the cell is classified to be in state $i$.

Transition probability from state $i$ to $j$ is given by

$$p_{ij} = \frac{n_{ij}}{\sum_{k \neq i} n_{ik}}, \tag{9}$$

where $n_{ij}$ is the number of transitions from state $i$ to state $j$.

Transition rate from state $i$ to $j$ is defined as

$$q_{ij} = \frac{p_{ij}}{\langle T_i \rangle}, \tag{10}$$

where $\langle T_i \rangle$ is the mean duration of state $i$.

The survival probabilities for state $i$ are defined by

$$P\left(T_i > \tau\right) = p_i \frac{N\left(T_i > \tau\right)}{N\left(T_i > 0\right)}, \tag{11}$$

where $N\left(T_i > \tau\right)$ is the number of instances where the duration of state $i$ is longer than $\tau$.

##### 9.4 Beat frequency during a run

The cilia beat frequency for each run period longer than 1 s was estimated using a fast Fourier transform analysis of the raw speed (i.e. calculated using the raw centroid positions). A second order Savitzky–Golay filter with frame length 45 was used to reduce the noise of the Fourier transform and the beat frequency for each run was taken to be the highest peak within 30-90 Hz (or if the highest peak was twice the frequency of the second highest peak, then the latter was taken as the beat frequency).
